# The Innate Immune Response to Ischemic Injury: a Multiscale Modeling Perspective

**DOI:** 10.1101/249599

**Authors:** Elena Dimitrova, Leslie A. Caromile, Reinhard Laubenbacher, Linda H. Shapiro

## Abstract

**Background:** Cell death as a result of ischemic injury triggers powerful mechanisms regulated by germline-encoded Pattern Recognition Receptors (PRRs) with shared specificity that recognize invading pathogens and endogenous ligands released from dying cells, and as such are essential to human health. Alternatively, dysregulation of these mechanisms contributes to extreme inflammation, deleterious tissue damage and impaired healing in various diseases. The Toll-like receptors (TLRs) are a prototypical family of PRRs that may be powerful anti-inflammatory targets if agents can be designed that antagonize their harmful effects while preserving host defense functions. This requires an understanding of the complex interactions and consequences of targeting the TLR-mediated pathways as well as technologies to analyze and interpret these, which will then allow the simulation of perturbations targeting specific pathway components, predict potential outcomes and identify safe and effective therapeutic targets.

**Results:** We constructed a multiscale mathematical model that spans the tissue and intracellular scales, and captures the consequences of targeting various regulatory components of injury-induced TLR4 signal transduction on potential pro-inflammatory or pro-healing outcomes. We applied known interactions to simulate how inactivation of specific regulatory nodes affects dynamics in the context of injury and to predict phenotypes of potential therapeutic interventions. We propose rules to link model behavior to qualitative estimates of pro-inflammatory signal activation, macrophage infiltration, production of reactive oxygen species and resolution. We tested the validity of the model by assessing its ability to reproduce published data not used in its construction.

**Conclusions:** These studies will enable us to form a conceptual framework focusing on TLR4-mediated ischemic repair to assess potential molecular targets that can be utilized therapeutically to improve efficacy and safety in treating ischemic/inflammatory injury.

## Introduction

Regardless of the initial insult, optimal healing of damaged tissue relies on the precise balance of pro-inflammatory and pro-healing processes of innate inflammation to the extent that variations in either arm can exacerbate many diseases from obesity to autoimmunity. Consequently, focusing on the mechanisms and molecules responsible for maintaining this delicate balance may identify novel regulatory nodes that are fundamental to the overall orchestration of tissue repair. Dissection of the steps by which these pivotal regulatory proteins operate will increase our understanding of these interdependent responses and allow the development of more specific, effective and clinically translatable therapeutic targets to enhance the healing process and improve clinical outcomes.

Tissue damage resulting from ischemic injury invariably leads to cell death and activates the same innate inflammatory responses triggered by pathogenic organisms. During inflammation, production of pro-inflammatory cytokines (initially by tissue resident macrophages) attracts monocytes from the blood stream to the injured tissue where they leave the vessels, accumulate, differentiate into macrophages and participate in the healing process by clearing the necrotic tissue and promoting tissue regeneration [3, 29–31], **Fig. 1).** The critical role of macrophages in post-ischemic healing is illustrated by studies in which systemic depletion of macrophages showed markedly impaired wound healing and perfusion recovery [32, 33]. Macrophages and other cells constitutively display members of germline-encoded Pattern Recognition Receptors (PRRs) that recognize molecular signatures shared by invading pathogens (Pathogen-associated molecular patterns, PAMPs) and endogenous ligands released from damaged cells (Danger-associated molecular patterns, DAMPs). Upon recognition of these distress signals, PRRs rapidly activate their associated cells to eradicate the infection, remove cell debris and heal the damage. Members of the Toll-like receptor (TLR) family are predominant PRRs expressed on the cell-surface or in endosomes that stimulate the precise signal transduction and gene expression programs that guide the innate immune response in response to PAMPs and DAMPs. Ten human and twelve murine TLRs have been identified and are differentially activated by different ligands. For example, TLR3 detects double-stranded viral RNA, while TLR4 specifically recognizes the PAMP lipopolysaccharide displayed by gram-negative bacteria. Importantly, TLR4 also recognizes a number of DAMPs released by damaged cells and thus is critical to proper healing following ischemic injury, such as myocardial infarction, peripheral artery occlusion, and stroke [36, 40–44]. Dysregulation of these pathways triggers what are often extreme inflammatory responses resulting in further tissue damage, prolonging and exacerbating the disease [28]. An intricate system of control points exists to ensure the proper response consisting of positive and negative regulators, feedback loops and cross-talk among signaling pathways.

**Figure 1.**
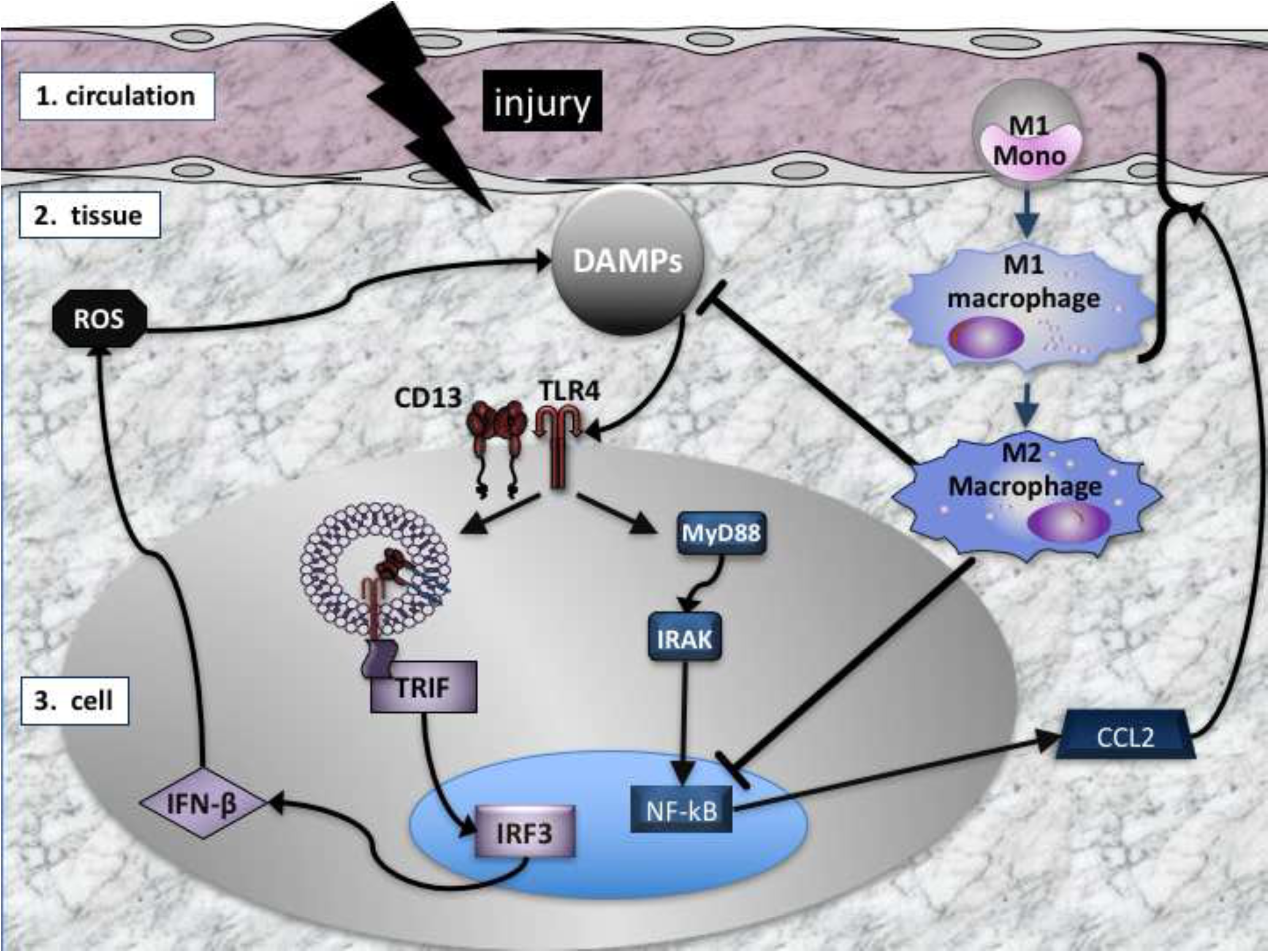
Scheme of the innate immune response to injury. Injury triggers the production of DAMPs in the tissue that activate intracellular responses via TLR4, initially on the resident macrophages (large gray oval). TLR4 activation stimulates two intracellular pathways, the MyD88 dependent (blue rectangles), resulting in production and secretion of the chemoattractant CCL2 which serves to recruit additional immune cells from the circulation (right). In response to CCL2, M1 monocytes leave the circulation and enter the tissue where they differentiate into pro-inflammatory M1 macrophages that clear toxic debris and become activated to produce more CCL2, perpetuating the inflammatory response. TLR4 can also signal via a MyD88-independent endocytic pathway (center left) that is mediated by CD13, TRIF and IRF3. Increased activation of this pathway can lead to production of cell-damaging ROS and increased DAMPs. Finally, M1 macrophages convert into pro-healing M2 macrophages which dampen the pro-inflammatory response by blocking production of CCL2 and DAMPs, leading to resolution.

Predicting and accurately testing the outcomes of targeting one or a combination of these nodes by biological methods is challenging, prompting us to create a mathematical model that captures the mechanisms involved at the tissue as well as cellular scale. This model then allows the simulation of interventions at either scale. As modeling framework we have chosen a time- and state-discrete model that captures the regulatory logic of the different mechanisms, and provides a qualitative description of model dynamics, without the need for quantitative kinetic and other parameters. Should it become necessary later to make quantitative assessments of processes, this discrete model can be converted into a continuous model with the same wiring diagram through the addition of parameters.

In recent years, a systems biology approach using mathematical modeling has been applied successfully to the study of events related to vascular injury. Several studies have focused on the molecular level, in particular the response of growth factors, such as VEGF [45–47]. In [48], the effect of ischemia/reperfusion on cardiomyocytes was explored through a mathematical model. Reperfusion-induced vasogenic edema was studied in [49] with the use of a mathematical model. There is evidence that nitric oxide can mitigate the negative effects of reperfusion, which is explored through mathematical modeling in [50, 51]. Other studies have focused on tissue-level phenomena such as hyperplasia formation or the effects of tissue oxygenation [52–55], or the mechanics of platelet deposition [56, 57]. The effect on the endothelial layer of blood vessels was modeled in [58]. A model of the innate and adaptive immune response to ischemic injury at the tissue level in the context of organ transplant surgery is presented in [59]. To our knowledge, no general mathematical models encompassing both the tissue and intracellular scales have been proposed for the innate immune response to vascular injury, making the model presented here novel.

## Results

### Description of the model

We created a network model based on numerous published biological studies of TLR4 signaling in response to injury or infection in the tissue (reviewed in [43, 44]) as well as our own studies of the role of CD13 in this response [16]. To capture the nature of the inflammatory response, we designed the model to initiate in the tissue (tissue scale) and release molecules which in turn trigger intracellular signaling mechanisms (cell scale), transcription and production of mediators that are secreted into the tissue to participate in a feedback loop to sustain further macrophage infiltration and wound healing. In the wiring diagram of the model (**Fig. 2**) injury is represented by the orange triangular node, which has two possible states, 0 and 1, indicating that injury is absent, respectively present. The production of DAMPs (purple circular node) can assume three possible states, representing ‘low, medium, high’, on the one hand, which impacts the cell scale by activation of signal transduction in resident macrophages (gray oval) on the other hand, which produces chemoattractants (CCL2) that recruit additional macrophages from the circulation (M1). Each resident or recruited macrophage responds to the presence of DAMPS by activating two pathways, resulting in the production and export of reactive oxygen species (ROS) and the inflammatory cytokine CCL2 (depicted as rectangular blue nodes in the model). ROS is considered as either present or absent, whereas CCL2 has three possible states, representing ‘low, intermediate, high.’ The M1 node in the tissue scale (black circular) can take on 3 states: with 0 representing the absence of macrophage activation; 1 representing the standard inflammatory response, initially as activation of resident macrophages or recruited macrophages as the response progresses; and 2 corresponding to the exaggerated recruitment of pro-inflammatory macrophages in exacerbated injury. As the healing process progresses, M1 macrophages differentiate into pro-healing M2 macrophages (purple circular M2 node) and, among other effects, influence the intracellular pathways in the monocytes and decrease DAMPs to diminish the pro-inflammatory response.

**Figure 2.**
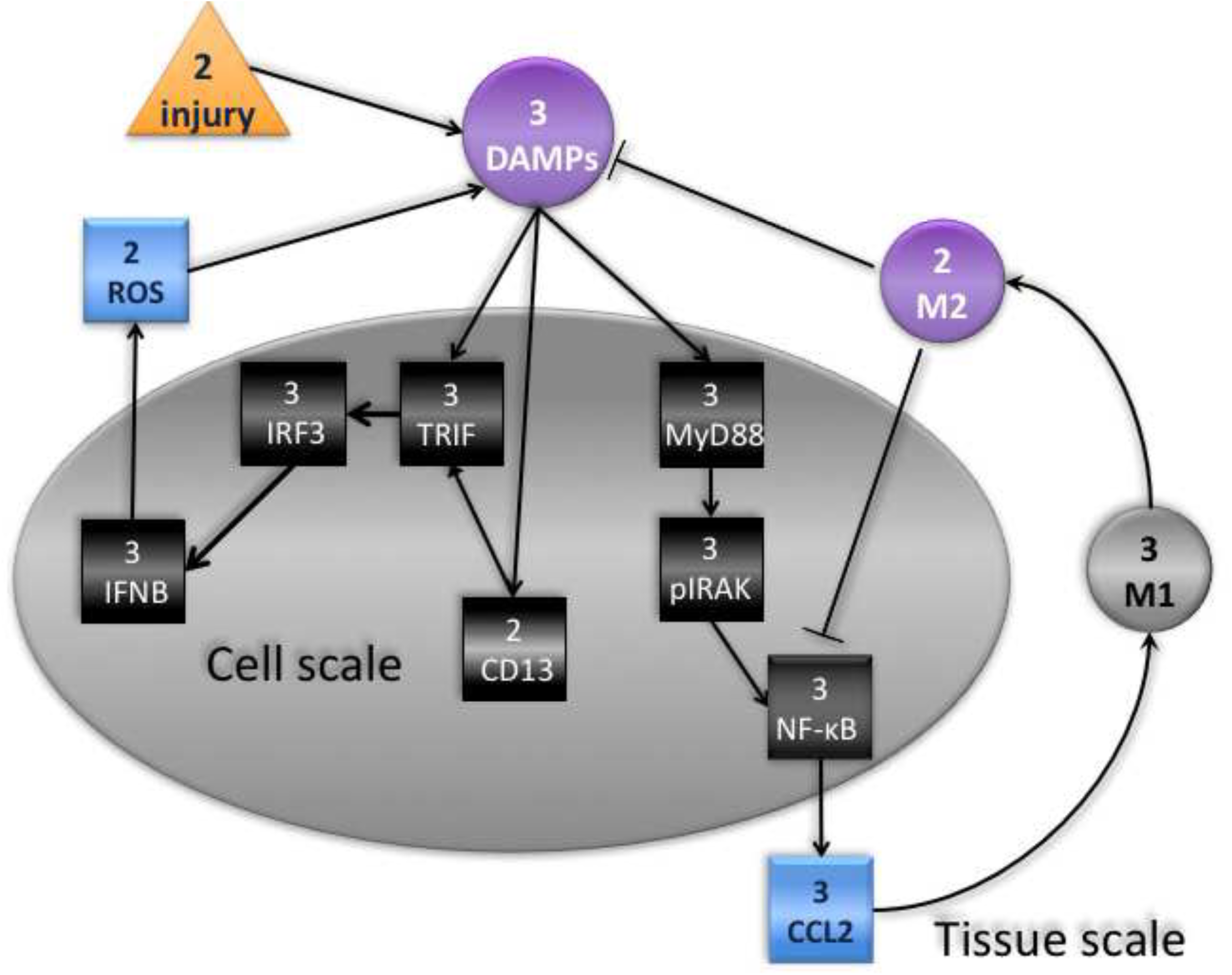
Wiring diagram of the model. Injury (orange triangle) has two possible states, 0- absent, and 1- present. The response to injury occurs at two simultaneous scales, the internal cell scale (gray oval) and the extracellular tissue scale. The tissue scale initiates with production of DAMPs (purple circle) with three states, low, medium, high, and the intracellular activation of resident macrophages via the MyD88-dependent (MyD88/IRAK/NF-κB/CCL2) and –independent (CD13/TRIF/IRF3/IFN-β) pathways, resulting in recruitment of additional immune cells from the circulation (M1) and/or production of toxic ROS. The M1 node (black circle) can take on 3 states: 0, absence of macrophage activation, including resting resident macrophages; 1 standard inflammatory response- initial activation of resident macrophages and later, of recruited macrophages; and 2 exaggerated recruitment of pro-inflammatory macrophages in exacerbated injury. As the process continues, M1 macrophages become pro-healing M2 macrophages (purple circle) and dampen the pro-inflammatory response.

While hundreds of intracellular and extracellular molecules have been connected to the TLR4 pathway, we have limited our nodes primarily to those with published knockout studies with the understanding that we will eventually expand upon this basic model. Finally, we have made numerous assumptions to simplify the model. Specifically, we have assumed that the degree of injury is such that there is a likely probability of resolution and that injury induces uniform responses at all levels regardless of individual attributes of the tissue, cell or molecules. Similarly, we have assumed that the response to injury is singularly mediated by the TLR4 pathway and that tissue resident macrophages only participate in the initiation of the response but not at later steps. We have narrowly restricted our nodes and response outcomes within this pathway to a defined set of effectors, omitting numerous others that have been implicated in this response. The most conspicuous example of this is TLR4 itself: *while we are modeling the TLR4-mediated response to injury, TLR4 is not a node in the model as it simply relays external signals to the cell interior*. These assumptions can be modified and elaborated upon as the model evolves.

### Biological mechanisms and translation into logical rules

### Description of the Model

**Table 1** contains a description of all the network nodes in the model, together with the possible states they can assume. The arrows in the diagram in **Fig. 2** represent the dependencies between network nodes, that is, all of the regulatory inputs that the node receives from other nodes. **Table 2** lists the logical rules that we have developed to translate our biological observations into qualitative effects on the different nodes. When applied to the various input node values, these rules will determine the state of the node at the next time step. They are grouped according to the scale at which they operate, with the tissue scale rules listed first. The effect of these rules on the state of a particular node can be captured through a “transition table”. **Table 3** is an example of the transition table for the node TRIF, which depends on DAMPs and CD13. All possible input configurations for DAMPs (0, 1, 2) and CD13 (0, 1) are specified in columns 1 and 2. By applying the rules, we can assign state values to TRIF (column 3) that would logically result from these input combinations.

**Table 1.** List of species, model states and biological characteristics

| Table 1. List of species, model states and biological characteristics |  |  |  | model states |  |  |
| --- | --- | --- | --- | --- | --- | --- |
| species | # states | class | type | 0 | 1 | 2 |
| Injury | 2 | external stimulus | effector | absent | present | ---- |
| DAMPs | 3 | protein | effector | no injury | intermediate | high |
| M1 | 3 | cell | promotes inflammation | low | intermediate | high |
| M2 | 2 | cell | promotes healing | low | high | ---- |
| CD13 | 2 | protein | regulator | inactive | active | ---- |
| TRIF | 3 | protein | adaptor | inactive | active | hyperactive |
| IRF3 | 3 | protein | transcription factor | inactive | active | hyperactive |
| IFNβ | 3 | protein | cytokine | low | intermediate | high |
| ROS | 2 | chemical | effector | low | high | ---- |
| MyD88 | 3 | protein | adaptor | inactive | active | hyperactive |
| pIRAK | 3 | protein | kinase | inactive | active | hyperactive |
| NF-kB | 3 | protein | transcription factor | inactive | active | hyperactive |
| CCL2 | 3 | protein | inflammatory cytokine | low | intermediate | high |

**Table 2.** Tissue Scale Rules

| Table 2. Tissue Scale Rules |  |  |  |
| --- | --- | --- | --- |
| Rule |  | Literature support | Relevant references |
| **CCL2 and ROS <- from cell model** |  |  |  |
| 1 | Injury (2) = 0 if M2 =1 and previous injury = 1 | M2 macrophages will resolve tissue damage due to injury. | [1-4] |
| 2 | DAMPs (3) = 0 if no Injury AND no ROS regardless of M2 | DAMPs are generally not accessible without tissue damage. | [5, 6] |
| 3 | DAMPs =0 if (Injury XOR ROS) and M2=1 | M2 macrophages can completely resolve damage due to either injury or ROS. | [5-12] |
| 4 | DAMPs =1 if (Injury XOR ROS) and M2=0 unless previous DAMPs = 2 | Lack of M2 macrophages leads to increased tissue damage in response to injury or ROS. | [5-12] |
| 5 | DAMPs =1 if (Injury AND ROS) and M2=1 | Extensive damage resulting from both injury and ROS in the presence of M2 is not completely resolved. | [5-12] |
| 6 | DAMPs =2 if (Injury AND ROS) and M2=0 | Excess injury triggers an overwhelming immune response that destroys the tissue in the absence of M2 macrophages. | [5-12] |
| 7 | M1 (3) = 0 if (CCL2 = 0) | Pro-inflammatory cytokines (exemplified by CCL2) are required to recruit M1 monocytes/macrophages. | [1-4, 13-15] |
| 8 | M1 = 1 if CCL2 = 1 | Macrophage recruitment is initiated in response to cytokines. | [1-4, 13-15] |
| 9 | M1 = 2 CCL2 = 2 | increased cytokine levels will recruit more M1 macrophages. | [1-4, 13-15] |
| 10 | M2 (2) = 1 if M1 =1 | M1 macrophages differentiate into M2. | [1-4, 13-15] |
| 11 | M2 = 0 otherwise | M2 macrophages are derived from M1; overwhelming M1 infiltration overcomes M2. | [1-4, 13-15] |
| **DAMPs and M2 -> to cell scale** |  |  |  |
| Cell Scale Rules |  |  |  |
| Rule |  | Literature support | Relevant references |
| **DAMPs and M2 <- from tissue model** |  |  |  |
| 12 | CD13 (2) = 1 if DAMPs = 1 or 2 | CD13 is phosphorylated upon ligand binding to TLR4 | [16, 25, 26] |
| 13 | CD13 = 0 otherwise | CD13 is not activated without inflammation | [16] |
| 14 | TRIF (3) = 0 if DAMPs = 0 regardless of CD13 | There is no response without tissue damage. | [16, 27, 28] |
| 15 | TRIF = 1 if (DAMPs = 1) and (CD13 = 1) | Ligation and endocytosis of TLR4 triggers TRIF activation. | [16, 27, 28] |
| 16 | TRIF = 2 if (DAMPs = 1) and (CD13 = 0) | TRIF is hyper-activated in the absence of CD13 | [16, 27, 28] |
| 17 | TRIF = 2 if DAMPs = 2 regardless of CD13 | Excess injury triggers an overwhelming immune response. | [16, 27, 28] |
| 18 | IRF3 (3) = TRIF (3) | TRIF activates IRF3 | [16, 27, 28] |
| 19 | IFN-β (3) = IRF3 | Active IRF3 transcriptionally activates IFN-β | [16, 18, 34-36] |
| 18 | ROS (2)= 1 IFNβ = 2 -> to cell model | High levels of IFN-β induce ROS | [7, 8, 10-12, 16] |
| 19 | ROS = 0 otherwise | Low levels of IFN-β do not induce ROS. | [7, 8, 10-12, 16] |
| 20 | MyD88 = DAMPs (3) | DAMPs bind TLR4 and activate MyD88 from the cell surface. | [37-39] |
| 21 | pIRAK = MyD88 (3) | Activated MyD88 enables IRAK phosphorylation/activation. | [37-39] |
| 22 | NF-κB = 0 if M2 = 1 and (pIRAK = 0 or 1) | M2 macrophages dampen NF-κB activity and halt inflammation unless overwhelming response. | [37-39] |
| 23 | NF-κB = pIRAK (3) otherwise | pIRAK activates NF-κB. | [14, 15] |
| 24 | CCL2 = NF-κB (3) | NF-κB transcriptionally regulates CCL2 | [37-39] |
| **CCL2 and ROS -> to tissue scale** |  |  |  |

**Table 3.** TRIF Depends on DAMPs and cd13

| DAMPs | CD13 | TRIF |
| --- | --- | --- |
| 0 | 0 | 0 |
| 0 | 1 | 0 |
| 1 | 0 | 2 |
| 1 | 1 | 1 |
| 2 | 0 | 2 |
| 2 | 1 | 1 |

### Initiation of the tissue scale: Injury, cell death and TLR4 activation

We have focused the model on macrophage recruitment and included two mechanisms by which products of the intracellular pathways attract these effector cells to the site of injury. Initially, in response to tissue injury, dead and dying cells release endogenous intracellular proteins, thus providing molecular ‘danger’ signals or DAMPs (Table 2, rules #2-6, refs. [36, 41]. The extracellular DAMPs activate tissue-resident macrophages [60] and trigger the intracellular signaling cascades of the inflammatory response that serve to initially recruit circulating macrophages to the site of injury to repair damaged tissue, remove dead cells and heal the wound. Paradoxically, failure to activate this response results in further damage due to inflammatory hyper-activation by the toxic accumulation of apoptotic cell debris, whereas excessive activation can also lead to dysregulated inflammation and further tissue damage. Therefore, tight control of the response to injury is imperative for a balanced and effective immune response.

### Intracellular signaling pathways from the plasma membrane and endosome

Once activated, the TLR4 response to DAMPs is somewhat unique in that it activates two distinct intracellular signaling pathways from different locations. These can be distinguished by their requirement for the intracellular adaptor protein MyD88. MyD88-dependent signaling originates from the plasma membrane, inducing the classic pro-inflammatory cascade [24, 61, 62]. Alternatively, MyD88-independent, TRIF-mediated signals originate from intracellular endosomal vesicles, activation of transcription and production of proteins that generally promote the adaptive immune response [24]. The importance of controlling these signaling pathways is illustrated by the induction of severe pathologies resulting from overstimulation of the pathway or the production of deleterious reactive oxygen species (ROS) by excessive levels of MyD88-independent signaling. ROS release into the tissue damages cells, increasing tissue DAMPs and amplifying the immune response. Finally, systemic depletion of macrophages severely impairs wound healing [22, 23], suggesting that independent but overlapping regulatory nodes exist [63].

### MyD88-dependent signal transduction from the plasma membrane

DAMPs recruit MyD88 to the plasma membrane to result in the phosphorylation of IRAK (Interleukin-1 receptor-associated kinase 1) to pIRAK, which then disassociates from MyD88 to perform a series of additional interactions leading to activation and nuclear localization of the NF-κB (nuclear factor kappa enhancer of B cells) transcription factor complex. In the nucleus, NF-κB induces the production of various inflammatory cytokines, such as CCL2, TNF-α, IL-12 and IL-1. We have chosen to focus on CCL2, but the other cytokines and their regulators can be added in the future. These factors are secreted from the cell to attract other inflammatory cells via their cognate receptors, ultimately impacting the tissue model by recruiting more macrophages, which can either facilitate healing in a balanced state or escalate tissue damage when dysregulated. The amplitude of these components is determined largely by the intensity of DAMPs. We have assigned three levels of activation to MyD88, IRAK, NF-κB and CCL2 (0, 1, 2) Table 2, Rules #20-24.

### MyD88-independent signal transduction from endosomes

Alternatively, ligand binding to TLR4 also induces translocation of TLR4/ligand from the plasma membrane into endosomal vesicles [64]. Positive and negative regulators of this process exist and represent additional nodes for future inclusion [65, 66]. This pathway involves the TRIF (TIR domain-containing adaptor protein-inducing IFN-β) adaptors to activate the interferon regulatory factors, IRFs, a family of transcription factors that are important in antiviral defense, cell growth and immune regulation. One of these, IRF3, stimulates production of the type I interferons, IFN-α and –β (designated as IFN-β). IFN binding to IFNAR (the IFN-α and -β receptor, not included as a node) induces signal transduction to initiate production of iNOS, the enzyme responsible for the formation of bactericidal reactive oxygen species (ROS). While the secreted extracellular ROS are critical to microbial defense, these can be toxic when present at high levels and lead to further tissue injury, cell death, increased release of DAMPs and recruitment of macrophages in the tissue [67–70]. The hyperactivated state of this pathway (IFN-β=2) triggers ROS, while normal response to injury produces IFN-β but no ROS. We have assigned three levels of activation to TRIF, IRF3, IFN-β (0, 1, 2) and two to ROS (0,1), **Table 2,** Rules #14-19. Finally, this pathway also triggers a distinct, delayed alternate pathway to NF-κB activation [62] which we have not included in this acute model.

### Tissue injury resolution or further damage

Cytokines produced intracellularly are secreted into the tissue where they activate endothelial cells lining adjacent blood vessels to attract additional pro-inflammatory M1 monocytes to enhance the response [71]. We assume in the model that these cytokines are initially produced by tissue-resident macrophages and subsequently by recruited, infiltrating M1 macrophages (**Table 2**, rules #7-9). Once in the tissue, M1 monocytes differentiate into M1 macrophages that ingest the DAMPs and degrade the extracellular matrix to allow development of granulation tissue and the eventual scar. Reduced DAMPs levels prompt a second, pro-resolution phase where M1 macrophages switch to an M2 phenotype (rules 10, 11, ref [15]). M2 macrophages contain fewer inflammatory molecules and proteases and elicit factors that promote angiogenesis and collagen deposition as well as reduce inflammation by downregulating intracellular NF-κB activity and CCL2 production (**Table 2**, rules #1, 22 and 24, ref. [15]). A systemic lack of monocytes/macrophages leads to persistence of DAMPs, increased overall cytotoxic TLR4 signaling, lack of M2 macrophages and further damage [72]. Similarly, a lack of M2 macrophages also leads to persistent DAMPs, excess inflammatory cytokines, damaging oxidative stress and ROS production [73]. (**Table 2,** rules #3-6)

### CD13 in TLR4 Signaling

We have demonstrated that a lack of CD13 increases TLR4 MyD88-independent signaling by virtue of its endocytic regulatory properties [16]. We have also shown that CD13 is phosphorylated upon ligand binding, which is required for its effects on receptor uptake [16, 25]. This rise in ligand-receptor internalization enhances activation of the MyD88-independent endosomal-signaling arm of the TLR4 response, leading to aberrantly high levels of type I interferons and ultimately production of injurious reactive oxygen species (ROS), thus exacerbating injury due to inflammation. We have incorporated results from this study into the model, where CD13 = 0 when unphosphorylated/inactive, or CD13 = 1 when phosphorylated/activated (**Table 2**, Rules #12-17).

### Model simulation

Below we describe the results of a model analysis and validation of the model by comparing its behavior under certain perturbations with known, previously published *in vivo* results from knockout animal studies (references listed in **Table 5**). Interrogation of the model is through simulation. The model is first initialized with all possible state values for each of the nodes, (e.g. Injury = 0, 1, DAMPs = 0, 1, 2, etc.). We then apply the rules in **Table 2** to each of the model nodes to obtain the new state value for each node according to our rules. Further iteration provides a chronological time course of states, which can either terminate in a steady state or a periodic repeated pattern or ‘limit cycle’. For our model, all time courses terminate in a steady state. However, since the model integrates two different spatial scales and consequently, two different temporal scales, we needed to modify the scheme by which the nodes are updated. Since we assume the intracellular scale will be significantly faster than the tissue scale, we have designed the update scheme as follows: for a given initialization for all nodes, we first combine the nodes from the cell model, the two input nodes DAMPs and M2 and the two output nodes ROS and CCL2 and together consider them as a separate model. We then iterate this sub-model until it reaches a steady state. The steady state values that are obtained for the two output nodes are assigned as initialization values for the tissue level nodes to enter into the rule simulation. The new values of the tissue-level nodes reached at the end of the simulation, merged with the steady state values of the cell model, then comprise the state of the entire model at the next time step.

### Model analysis

The initial model analysis below was obtained by exhaustively simulating the model by computing the transition for each possible configuration of node values, using the software package PlantSimLab (http://app.plantsimlab.org). In this way we can determine all possible steady states of the model, which in this case can be interpreted as all the possible outcomes of the response to injury, taking into account all possible configurations of the underlying network. For each steady state, we record the number of states that eventually lead to a particular steady state/outcome, also known as the ‘basin of attraction’ of this steady state. This provides a measure of how likely the different outcomes are. For clarity, we have listed the outcomes for the intracellular and tissue components of the model separately **(Tables 4 and 5)**. Simulations of the cell-level model from all initial states with DAMPs and M2 as input nodes results in six possible steady states, all having the same sized basin of attraction, that reflect the response to various input levels of DAMPs **(Table 4)**. Components 1 and 2 portray the response in the cell where there is either no injury or injury has been resolved. Components 3 and 4 describe the chronic response to initial injury and finally, Components 5 and 6 describe the states where high levels of cytokines and ROS lead to cell death, overwhelming the inflammatory response. Values generated by the cell model initiate the tissue model with the input nodes CCL2 and ROS **(Table 5).** This simulation results in a major steady state component 1 (92.6%) that describes the tissue with low levels of DAMPs and macrophages as would result with either no injury or injury followed by resolution. In comparison, Component 2 is a state with a small basin of attraction (5.6%), that is, a steady state observed rarely, that represents an overwhelming inflammatory response triggered by injury with high levels of cytokine production, ROS and cell death, as demonstrated by maximal levels of all pro-inflammatory components and ROS. Finally, since the simulation software initializes from all possible values, it can produce biologically improbable steady states as in Component 3 where ROS is present with no injury. This is reflected by the fact that the basin of attraction for this steady state only contains less than 2% of all possible model initializations.

**Table 4.** Cell Scale Steady States

| | DAMPs(3) | M2(2) | ROS(2) | CCL2(3) | TRIF(3) | CD13(2) | IRF3(3) | INFb(3) | MyD88(3) | pIRAK(3) | NF-κB(3) | basin of attraction |
| --- | --- | --- | --- | --- | --- | --- | --- | --- | --- | --- | --- | --- |
| Component 1 | 0 | 0 | 0 | 0 | 0 | 0 | 0 | 0 | 0 | 0 | 0 | 16.66% |
| Component 2 | 0 | 1 | 0 | 0 | 0 | 0 | 0 | 0 | 0 | 0 | 0 | 16.66% |
| Component 3 | 1 | 0 | 0 | 1 | 1 | 1 | 1 | 1 | 1 | 1 | 1 | 16.66% |
| Component 4 | 1 | 1 | 0 | 0 | 1 | 1 | 1 | 1 | 1 | 1 | 0 | 16.66% |
| Component 5 | 2 | 0 | 1 | 2 | 2 | 1 | 2 | 2 | 2 | 2 | 2 | 16.66% |
| Component 6 | 2 | 1 | 1 | 2 | 2 | 1 | 2 | 2 | 2 | 2 | 2 | 16.66% |

**Table 5.** Tissue scale steady states

|  | DAMPs(3) | M1(3) | M2(2) | Injury(2) | ROS(2) | CCL2(3) | basin of attraction |
| --- | --- | --- | --- | --- | --- | --- | --- |
| Component 1 | 0 | 0 | 0 | 0 | 0 | 0 | 92.59% |
| Component 2 | 2 | 2 | 0 | 1 | 1 | 2 | 5.55% |
| Component 3 | 2 | 2 | 0 | 0 | 1 | 2 | 1.85% |

### Model validation

To verify that the model captures some key features of the injury response, we considered published studies of injury models in wild type animals and those engineered to lack one of five different nodes in our model and interpreted the phenotypes in light of our model behavior [3, 16–24, 74, 75]. Similar to simulations of the wild type models, we initially computed the steady state values for each intracellular component from all possible initializations with the specific node knocked out (essentially set to 0), represented by the numbers in each row **(Table 6)**. These intracellular steady state values were then assigned as input values to initialize the tissue model and then we determined the values at which the output converged (the steady state) as described below.

**Table 6.**
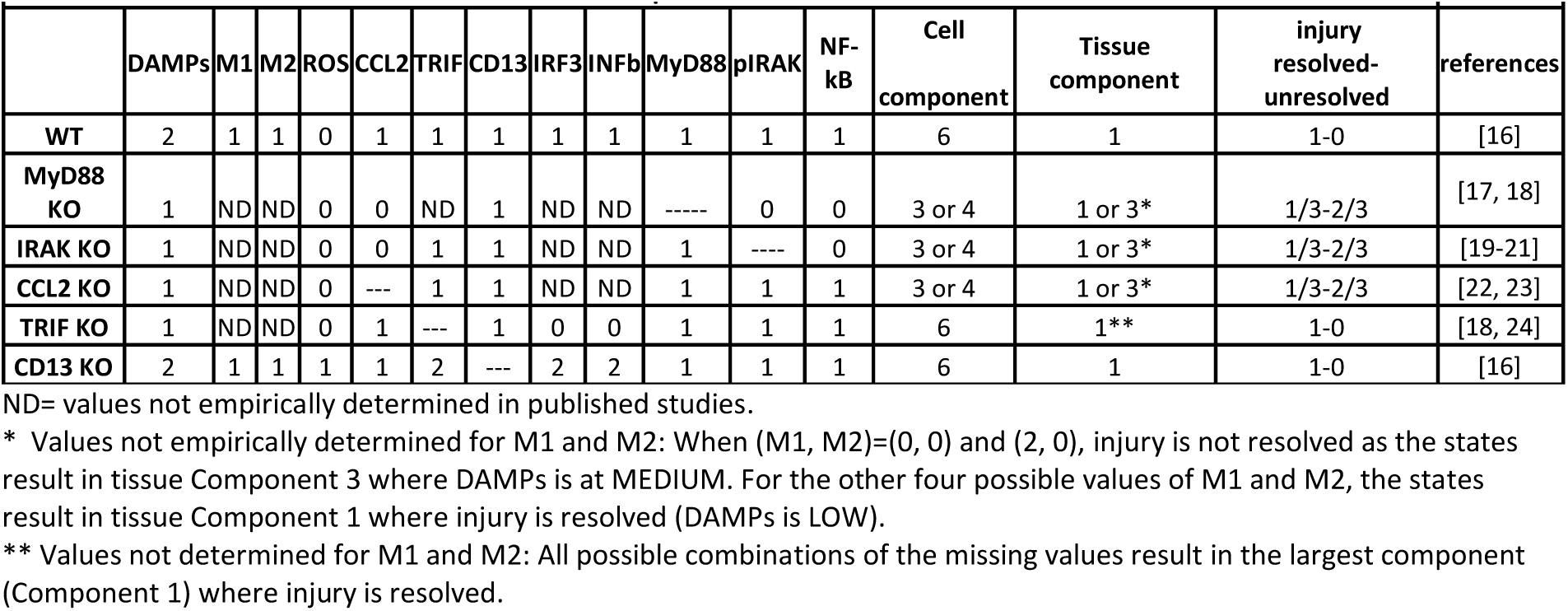
Verification of the model based on results from published studies.

|  | DAMPs | M1 | M2 | ROS | CCL2 | TRIF | CD13 | IRF3 | INFb | MyD88 | pIRAK | NF-kB | Cell component | Tissue component | injury resolved-unresolved | references |
| --- | --- | --- | --- | --- | --- | --- | --- | --- | --- | --- | --- | --- | --- | --- | --- | --- |
| WT | 2 | 1 | 1 | 0 | 1 | 1 | 1 | 1 | 1 | 1 | 1 | 1 | 6 | 1 | 1-0 | [16] |
| MyD88 KO | 1 | ND | ND | 0 | 0 | ND | 1 | ND | ND | ----- | 0 | 0 | 3 or 4 | 1 or 3* | 1/3-2/3 | [17, 18] |
| IRAK KO | 1 | ND | ND | 0 | 0 | 1 | 1 | ND | ND | 1 | ---- | 0 | 3 or 4 | 1 or 3* | 1/3-2/3 | [19-21] |
| CCL2 KO | 1 | ND | ND | 0 | --- | 1 | 1 | ND | ND | 1 | 1 | 1 | 3 or 4 | 1 or 3* | 1/3-2/3 | [22, 23] |
| TRIF KO | 1 | ND | ND | 0 | 1 | --- | 1 | 0 | 0 | 1 | 1 | 1 | 6 | 1** | 1-0 | [18, 24] |
| CD13 KO | 2 | 1 | 1 | 1 | 1 | 2 | --- | 2 | 2 | 1 | 1 | 1 | 6 | 1 | 1-0 | [16] |

ND= values not empirically determined in published studies.
\* Values not empirically determined for M1 and M2: When (M1, M2)=(0, 0) and (2, 0), injury is not resolved as the states result in tissue Component 3 where DAMPs is at MEDIUM. For the other four possible values of M1 and M2, the states result in tissue Component 1 where injury is resolved (DAMPs is LOW).
\*\* Values not determined for M1 and M2: All possible combinations of the missing values result in the largest component (Component 1) where injury is resolved.

## Intracellular states

### Wild type, TRIF knockout, and CD13 knockout

The states in each of these simulations converge to steady states in the intracellular model which correspond to states in the tissue model that proceed to resolution (Component 1), in agreement with the tissue states.

### MyD88, IRAK, and CCL2 knockouts

The intracellular states again lack input values for M2 and so both possible values, LOW and HIGH, are considered. When M2 = LOW, the given state is a steady state itself and when input into the tissue model, it converges to the steady state of Component 3, where injury fails to resolve as resulted from the absence of M2. This is precisely what happens for two of the six states of the tissue state. In the case where we set M2 = HIGH, then the simulation values correspond to injury resolution in the tissue.

## Tissue states

### Wild type and CD13 knockout

The experimental state is in the largest component (Component 1) generated by the model simulation, where injury is resolved.

### MyD88, IRAK, and CCL2 knockouts

The published studies did not determine values for a number of the states, particularly differentiating between M1 and M2. If we initialize the model with the two possible combinations of the missing values, (M1, M2) = (0, 0) and (2, 0), injury is not resolved as DAMPs is at MEDIUM level (Component 3 in the simulation), suggesting that the injury will eventually resolve unless M2 macrophages are absent, or 0. For the other possible values of M1 and M2, (0, 1), (1,1), (0,0) and (2,1), the states are in the largest component (Component 1) where injury is eventually resolved.

### TRIF knockout

The values for M1 and M2 are again missing but all possible combinations of values give states that are in the largest component (Component 1), where injury is resolved.

Taken together, the model we have constructed essentially resolves the injury despite perturbation with the exception of the absence of M2 macrophages. Since M2 cells are derived from M1 macrophages, the scenario where M1 is assigned as 0 and M2 as 1 is biologically impossible. Therefore, it can be assumed that the absence of M1 macrophages will also be considered to result in failure to resolve injury.

### Reconciliation with published studies

While we consider the results of the simulation to be consistent with the known experimental results, we are aware that states in **Table 6** do not necessarily match the published results of the *in vivo* experiments, but rather represent the steady states to which these biological systems would be expected to eventually converge. For example, experiments evaluating the response at 3-5d post injury during the inflammatory phase in the absence of the MyD88-dependent pathway generally report reduced inflammation [59, 60]. By contrast, interruption of the MyD88-independent pathway injury produces a pro-inflammatory, high damage state despite the absence of ROS, suggesting that the MyD88-dependent pathway contributes to inflammation-induced damage to a greater extent than the MyD88-independent pathway [18]. However, these experimental measurements are not taken at the point of equilibrium, but at defined time points (days post-injury) where the system is actively working toward resolving the injury. Therefore this is not a shortcoming of the model, but confirms that the model captures the most crucial features of the biological system.

### Summary and future directions

We have constructed a basic model of inflammatory signaling and monocyte trafficking in response to acute, sterile tissue injury that faithfully recapitulates components of published *in vivo* knockout experiments. Reconciling computational models with experimental data is difficult for a number of reasons. Biologists perturb systems with the goal of determining the intermediate steps that the system undergoes to achieve the steady state, in this case healing. Therefore we collect data on defined nodes at various time intervals following initiation of the experiment and rarely at a steady state. On the other hand, computational models test every possible combination of input values and converge on a steady state that can be considered as the long-term outcome of tissue injury. In the case of the fully functioning system in wild type animals, the damage is eventually resolved, and the intermediate steps proceed to the steady state of healing. In the case of loss of one of the nodes of the system, the model is perturbed, but eventually converges to resolution.

To this point, we have not modeled fibrosis and scarring which are often exacerbated when inflammation is dysregulated and can severely impact functional recovery of the tissue following ischemic injury. Including these processes in the model would likely capture the impairment of tissue function that persists following the resolution of inflammation in a compromised host.

We developed the current model as a basis for constructing a larger, more complex network model that can be used to predict the inflammatory response to different stimuli, additional receptors, cytokines, control points and cell types. For example, while we have included CD13 as a negative regulator of the MyD88-independent response, additional control nodes such as ATF3 (induces a negative feedback loop [65]) or the positive regulator CD14 (required for MyD88-independent signaling) could be added [66]. Alternatively, a component of gram negative bacterial cell walls triggers the same responses that we have modeled in response to injury. However, recurrent bacterial infections produce antibodies that bind to the bacteria, thereby creating a dual stimulus for the cell (via TLR4 and FcRs) to elicit a combined immune response considerably different from that initiated by either receptor alone and more efficient at triggering both innate and adaptive immunity [76, 77]. Mathematical modeling of such altered responses could lead to the identification of novel convergence nodes as therapeutic targets for inflammatory and autoimmune diseases.

A significant limitation of the current model is that it does not account for the fact that conditions in the tissue are not homogeneous so that the inputs to the intracellular component of the model vary across the tissue. In further work, we plan to construct a spatially heterogeneous model for the tissue scale, and each monocyte agent is equipped with its own intracellular network that can respond properly to local tissue conditions [78].

## Declarations Statements

- Ethics approval and consent to participate- Not applicable
- Consent to publish- Not applicable
- Availability of data and materials- Not applicable
- Competing interests- no competing interests
- Funding- NHLBI R01 127449 to LHS
- Authors’ Contributions- ED, LAC, RL and LHS participated in formulating the model, ED performed simulations, LAC and LHS provided validation data, ED, LHS and RL wrote the manuscript.
- Acknowledgements- We thank Charan Devarakonda and Mallika Ghosh for helpful discussions.

